# Effect Size and Power in fMRI Group Analysis

**DOI:** 10.1101/295048

**Authors:** Stephan Geuter, Guanghao Qi, Robert C. Welsh, Tor D. Wager, Martin A. Lindquist

## Abstract

Multi-subject functional magnetic resonance imaging (fMRI) analysis is often concerned with determining whether there exists a significant population-wide ‘activation’ in a comparison between two or more conditions. Typically this is assessed by testing the average value of a contrast of parameter estimates (COPE) against zero in a general linear model (GLM) analysis. In this work we investigate several aspects of this type of analysis. First, we study the effects of sample size on the sensitivity and reliability of the group analysis, allowing us to evaluate the ability of small sampled studies to effectively capture population-level effects of interest. Second, we assess the difference in sensitivity and reliability when using volumetric or surface based data. Third, we investigate potential biases in estimating effect sizes as a function of sample size. To perform this analysis we utilize the task-based fMRI data from the 500-subject release from the Human Connectome Project (HCP). We treat the complete collection of subjects (*N* = 491) as our population of interest, and perform a single-subject analysis on each subject in the population. We investigate the ability to recover population level effects using a subset of the population and standard analytical techniques. Our study shows that sample sizes of 40 are generally able to detect regions with high effect sizes (Cohen’s *d >* 0.8), while sample sizes closer to 80 are required to reliably recover regions with medium effect sizes (0.5 < d < 0.8). We find little difference in results when using volumetric or surface based data with respect to standard mass-univariate group analysis. Finally, we conclude that special care is needed when estimating effect sizes, particularly for small sample sizes.

## 1 Introduction

The analysis of multi-subject functional magnetic resonance imaging (fMRI) data is a critical component to most human brain mapping studies (Geuter et al., 2017). First, they provide an increase of the sensitivity of the overall experiment, as more data is available. Second, they allow one to determine whether the observed effects are common and stable across, or between, groups. Third, they allow for generalization of conclusions to the whole population of subjects.

Standard group analyses in the neuroimaging community typically involve utilizing two separate models. A first-level General Linear Model (GLM) analysis is performed on each subject’s data, which provides within-subject contrasts across parameter estimates (COPEs; e.g., activity magnitude estimates for [visual stimulation vs. rest]). A second-level analysis provides population inference on whether COPEs are significantly different from zero and assesses the effects of second-level predictors (e.g., group status, behavioral performance; Lindquist et al. (2012)). While, mixed-effects implementations exist for popular software packages (e.g., Woolrich et al. (2004); Friston et al. (2005)), researchers typically model first-level contrast data using an un-weighted, ordinary least squares (OLS) analysis (Mumford and Nichols, 2009).

Most group fMRI studies are small and have traditionally consisted of between 10-30 subjects (Button et al., 2013). Because of the relatively small number of subjects included, the sensitivity and reliability of these analyses are questionable and hard to investigate. In recent years there has been a great deal of discussion regarding how studies with small sample sizes undermine the reliability of neuroscience research (Button et al., 2013; MunafÒ et al., 2014). The argument is that small-sample studies have very low statistical power to detect true effects and a reduced likelihood for statistically significant results to be true effects. This has led to a situation where a large number of published results are not replicable and very likely false. Hence, a critical evaluation of the methods used to assess significance in small sample data is needed.

Traditional fMRI data consists of a time series of three-dimensional brain volumes, each composed of hundreds of thousands of equally sized voxels. Recently, cortical surface fMRI has experienced a rise in popularity (Fischl, 2012; Glasser et al., 2013). Here the cortical gray matter is represented as a 2-dimensional manifold surface in the format of a triangular mesh, which consists of vertices rather than voxels. This offers several potential advantages over volumetric fMRI data, as (i) it may improve alignment of cortical areas across multiple subjects, which is key for spatially accurate group-level analyses; and (ii) nearby locations are close in terms of distance along the cortical surface, and therefore tend to exhibit similar patterns of neuronal activity, while in volumetric fMRI locations that are close in terms of Euclidean distance may be from different areas of the cortex due to cortical folding or even from different tissue types. It remains an outstanding question whether surface based data is preferable, in terms of sensitivity and specificity, to volumetric data when performing standard group analysis.

After performing group analyses, it is often of interest to estimate and report effect sizes over certain key brain regions. Nearly a decade ago, a heated discussion about ‘circular’ or ‘nonindependent’ analysis in neuroimaging emerged in the literature (Kriegeskorte et al., 2010; Vul et al., 2009). Here an analysis is considered circular if it is based on data that were selected for showing the effect of interest or a related effect. Effect sizes estimated in a circular fashion tend to be significantly inflated compared to those estimated in a non-circular fashion. However, the size of these biases and how they depend upon sample size have not been further investigated to the best of our knowledge.

In this work, we first evaluate the effects of sample size on the sensitivity and reliability of the OLS approach, allowing us to evaluate the ability of small sampled studies to effectively capture population-level effects of interest. Second, we assess the difference between volumetric and surface based analysis in terms of their performance in group analysis in the mass-univariate setting where separate tests are performed at each brain location. Third, we investigate effect size distributions and potential biases in estimating effect sizes from real fMRI data (Kriegeskorte et al., 2010; Vul et al., 2009).

To explore these issues we utilize the task-based fMRI data from the 500-subject release from the NIH-sponsored Human Connectome Project (HCP) (Van Essen et al., 2013) for which processed volumetric and surface data are both available. We treat the complete collection of subjects (*N* = 491) as our population of interest, and perform single-subject analysis on each subject in the population. In particular, we focus our attention on fMRI from four different HCP tasks: emotion, gambling, motor, and working memory. This allows us to evaluate the appropriateness of the population-level assumptions required to perform valid second level analysis using a smaller sub-set of subjects from the population. Using this population of interest, we investigate how well we can estimate population-level effects using a smaller sample from the population, and common analytical techniques.

The proposed study shares similarities to previous work by Thirion et al. (2007), who used a cohort of 80 subjects to investigate the role inter-subject variability plays in sensitivity and reliability of studies with a smaller number (e.g., 10 - 16) of subjects. They showed, among other things, that 20 subjects or more should be included in functional neuroimaging studies in order for results to be replicable across samples (studies). Desmond and Glover (2002) recommended similar sample sizes of 20-30 subjects. Here we focus our analysis on a larger data set consisting of task-based fMRI data from *N* = 491 subjects. This allows us to investigate larger sample sizes than those in Thirion et al. (2007) and Desmond and Glover (2002), more variable tasks, and data sampled at a higher spatial and temporal resolution.

The task HCP data set has been used in several recent papers to investigate related topics. For example, Poldrack et al. (2017) used the data to obtain effect-size estimates for a number of common neuroimaging experimental paradigms in certain regions of interest. More recently, Lohmann et al. (2017) used it to investigate the false negative rates in studies with small sample sizes. Similar in spirit to our approach, they used a large cohort of 400 subjects to approximate the true positive effects. Thereafter, they computed the fraction of those effects that was detected reliably using standard software packages at various smaller sample sizes. They found that for common sample sizes this value was below 25%. In this work, we validate and expand upon the results from these papers.

This paper is organized as follows. We begin by focusing on the characteristics of the entire population of subjects. This allows us to perform a census to assess complete information regarding activation and variability within the population. We assess effect sizes across the brain and use this to create whole-brain maps of power and minimum sample sizes needed to detect certain effects. Next, we evaluate the effects of sub-sampling from the population, and evaluate how the reliability, power, and effect size of an fMRI study depends upon sample size. We conduct these analyzes on both the volumetric and surface based (grayordinate) data formats to compare them in terms of the ability to recover true effects. We conclude by comparing effect size estimates based on significant clusters, as they may be collected in pilot studies for the purpose of power analysis, against the population effects sizes.

## 2 Data and methods

### 2.1 Data sets

The Human Connectome Project 500 (HCP 500) consists of both structural and functional data from approximately 500 subjects. The functional data include both resting state (rfMRI) and task-related (tfMRI) images of multiple tasks. Our work here only uses the tfMRI GLM contrast images provided by the HCP. The following contrasts are considered: 2-back versus 0-back for the working memory (WM) task (*N* = 481), average motor action versus baseline for motor (*N* = 479), faces versus shapes for emotion (*N* = 471), and reward versus punishment for gambling (*N* = 479). The number of subjects for each task is slightly different since some subjects are missing a subset of the tasks. We also excluded subjects for which issues with either the segmentation or the surface quality control are reported (Issue code B) by the HCP consortium^1^

All data were acquired on a Siemens Skyra 3T scanner housed at Washington University in St. Louis. For each task, two runs were acquired, one with a right-to-left and the other with a left-to-right phase encoding. Whole-brain EPI acquisitions were acquired with a 32 channel head coil with TR= 720ms, TE= 33.1ms, flip angle= 52°, BW= *2290Hz/Px,* in-plane FOV = 208 × 180mm, 72 slices, 2.0mm isotropic voxels, with a multi-band acceleration factor of 8. For a full description of the fMRI Data Acquisition see Van Essen et al. (2012).

Volumetric imaging data was preprocessed according to the HCP ‘fMRIVolume’ pipeline (Glasser et al., 2013), which includes gradient unwarping, motion correction, fieldmap-based EPI distortion correction, brain-boundary-based registration of EPI to structural T1-weighted scan, non-linear registration into MNI152 space, grand-mean intensity normalization, and spatial smoothing using a Gaussian kernel with a FWHM of 4mm. Volumetric data were masked with a graymatter mask including cortical and subcortical graymatter to ensure that analyses for both data formats were conducted on comparable tissues.

Surface, or grayordinate, data was preprocessed according to the HCP ‘fM-RISurface’ pipeline (Glasser et al., 2013). Here, functional volume data from cortical gray matter voxels are mapped onto the standard 32k Conte69 surface mesh. The projection onto the standard surface mesh is accomplished via intermediate projections to each subject’s individual cortical surface and a high-resolution standard surface mesh (Glasser et al., 2013). Subcortical gray matter voxels are mapped onto the volumetric portion of the Conte69 space. The gray-ordinate data thus include data from cortical and subcortical graymatter in a standard space. Data were spatially smoothed using a Gaussian kernel with a FWHM of 4mm. For a full description of the pre-processing see (Glasser et al., 2013).

Analysis was performed using a general linear model (GLM) on the COPEs provided by the HCP in the S500 releasese. For each task, predictors (described below for each task) were convolved with a canonical hemodynamic response function to generate regressors. To compensate for slice-timing differences and variability in the HRF delay across regions, temporal derivatives were included and treated as variables of no interest. Both the data and the design matrix were temporally filtered using a linear high-pass filter (cutoff 200s). Finally, the time series was pre-whitened to correct for autocorrelation in the fMRI data.

Each of the datasets used in this paper are described in greater detail in Barch et al. (2013). Below follows a brief description.

#### 2.1.1 Working Memory

This task consisted of a version of the N-back task to assess working memory. Within each run, four different stimulus types (faces, places, tools and body parts) are presented in separate blocks. Half of the blocks use a 2-back task and the other half a 0-back task. A short (2.5s) cue indicates the task type at the start of the block. Each of the two runs contains 8 blocks consisting of 10 trials (2s stimulus presentation, 500ms ITI) and 4 fixation blocks (15s each). Each block contains 2 targets, and 2-3 non-target lures.

Eight task-related predictors were included in the GLM design matrix, one for each stimulus type in each of the N-back conditions. Each covered the period from the onset of the cue to the offset of the final trial. A linear COPE comparing 2-back versus 0-back was used for further analysis.

#### 2.1.2 Motor

In this task participants were presented with visual cues that ask them to tap their left or right fingers, squeeze their left or right toes, or move their tongue. Each block corresponds to one of the five movements and lasted 12s, and was preceded by a 3s cue. In each of the two runs, there are 13 blocks, with 2 tongue movements, 2 of each hand movement, 2 of each foot movement, and three 15s fixation blocks.

Five task-related predictors were included in the design matrix for each movement, each covering the duration of the 10 movements trials (12s). The cue was modeled separately. A linear COPE comparing the average of the different movements vs baseline was used for further analysis.

#### 2.1.3 Emotion Processing

In this task participants were presented with blocks of trials that either asked them to decide which of two faces shown at the bottom of the screen matched the face at the top, or which of two shapes presented at the bottom matched the shape at the top. The faces had either angry or fearful expressions. Trials are presented in blocks of 6 trials (2s stimulus presentation, 1s ITI) of the same task (‘face’ or ‘shape’). Each block was preceded by a 3s task cue (‘shape’ or ‘face’). Each of the two runs includes 3 face blocks and 3 shape blocks.

Two different task-related predictors were included in the design matrix, one corresponding to emotional faces and the other to the shape control condition. Each predictor covered a 21s duration composed of a cue and six trials. A linear COPE comparing emotional faces vs control was used for further analysis.

#### 2.1.4 Gambling

In this task participants were asked to guess the number on a mystery card in order to win or lose money. They are told that the number ranged from 1-9, and to guess whether the number on the mystery card number is greater or less than 5 by pressing one of two buttons. Feedback is the number on the card and either a green up arrow with $1 for reward trials; a red down arrow with $0.50 for loss trials; or the number 5 and a gray double headed arrow for neutral trials. The task is presented in blocks of 8 trials with mostly reward (6 reward trials —interleaved with either 1 neutral and 1 loss trial, 2 neutral trials, or 2 loss trials) or mostly loss (6 loss trials interleaved with either 1 neutral and 1 reward trial, 2 neutral trials, or 2 reward trials). In each of the two runs, there are 2 mostly reward and 2 mostly loss blocks, interleaved with 4 fixation blocks (15s each).

Two task-related predictors were included in the design matrix to model mostly reward and mostly punishment or loss blocks, each covering the duration of 8 trials (28s). A linear COPE comparing reward vs punishment blocks for gambling was used for further analysis.

### 2.2 Population Effect Size and Power

Throughout we assume that the population of interest consists of the full collection of roughly 500 subjects. For each task this allows us to compute population effect sizes at each voxel (or vertex for surface based results), based on all of the subjects in the population. This can subsequently be used for power analysis and for direct comparison with results obtained using data from a smaller subset of the population. This will allow for the evaluation of the influence of sample size on the results of group analysis.

There are multiple ways to compute effect size. Here we use a variant of Cohen’s d that can be computed at voxel v as the mean COPE (across subjects) divided by the standard deviation, i.e. *Ө_v_* = *μ*_*v*_/σ_*v*_. Using these values, we group voxels into four different effect size categories. Voxels are placed in the ‘low’ group if 0.2 ≤ │*θ*_*v*_│ < 0.5, the ‘medium’ group if 0.5 ≤ │*θ_v_*│< 0.8, and the ‘high’ group if 0.8 ≤ │*θ*_*v*_│. These groupings are based on guidelines from Cohen (1988). All remaining voxel are placed in a ‘no effect’ group.

The population effect sizes are used to construct ‘effect-size maps’ for each task, which are the effect sizes mapped onto their corresponding voxel or surface location. In addition, ‘power maps’ can be constructed, which are spatial maps of the estimated sample size needed to obtain p% power where *p* = 50, 80, 90, 95. Here power is computed based on a one-sided one-sample t-test with a = 0.001 and effect size θ_*v*_. These maps will allow us to identify the sample size required to detect an effect in a specific region with high power.

### 2.3 Effect of Sample Size on Group Analysis

In the continuation, we consider the 500 subject release population-level results as our ‘gold standard’, and use these results as a stand in for the ground truth effect size at different voxels for the various tasks. To evaluate the reproducibility of small sample fMRI studies, we draw *K* = 100 different samples from the population of each of the following sizes: *N* = 10, 20, 40, 60, 80 and 100. This allows us to mimic the typical sample sizes used in studies performed by individual labs. Note that throughout we are sampling with replacement as is performed in the Bootstrap procedure (Efron and Tibshirani, 1993). Hence, certain subjects may appear multiple times in any given sample. For each sample we perform a group analysis using the standard OLS approach. A voxel-wise threshold of p < 0.001 uncorrected is applied to the resulting t-maps to assess significance.

For each sample size we compute the proportion of studies in which a particular voxel or surface vertex is deemed active. To illustrate, begin by averaging all *K* binary activation maps (post threshold) obtained using a given sample size *N*. Next, denote the p-value of voxel v from the *k^th^* sample of size *N* by *p_v_,k,N*. Then, the proportion of studies in which voxel v is deemed activate for sample size *N* is given by

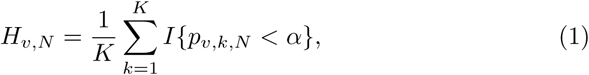

where *α* is the threshold and *I*{•} is the indicator function.

Next, we study how a voxel effect size corresponds to its likelihood to be deemed active. Using the population effect sizes groups defined in the previous section, we measured the proportion of voxels in the *K* samples that were deemed significant, as well as the proportion of voxels in each group that were active in more than 80% of samples. This allowed us to assess the false negative rate (under the assumption that voxels with this effect size should be deemed active) associated with each sample size and effect size, as well as the power to detect activation of a certain effect size.

### 2.4 Effects of Circular Analysis on Effect Size Estimation

In order to estimate the necessary sample size for an fMRI experiment when designing a study or writing a grant proposal, it is often recommended to carry out a small pilot study. Based on this pilot study one can then estimate the expected fMRI effect sizes for this specific task and setting. This might be done based on the significant activations or peak COPEs in the pilot study. Although many researchers are aware of the problems of a circular analysis that estimates an effect size based on a biased selection criterion (Kriegeskorte et al., 2010; Vul et al., 2009), it is unknown how much a circular analysis will actually bias the obtained estimates. Here, we compute effect sizes estimated averaged within significant clusters of each of the *K* samples of size *N*. We compare these estimates against the population effect sizes to estimate the bias introduced by the circularity.

## 3 Results

### 3.1 Population Effect Size and Power

We first characterized the distributions of COPEs and effect sizes (Cohen’s d) using the 500 subject population. Figure 1A shows the standard deviation of the COPEs across the approximately 500 subjects plotted against the mean COPE for each individual voxel. For all tasks, the standard deviation increases with the mean COPE, resulting in funnel shaped plots. The whole brain means of the COPEs are all positive (M = 2.13 for WM, M = 3.42 for motor, M = 5.12 for emotion, and M = 3.28 for gambling) as might be expected for an activation-hypothesis-based COPE. The gray and black lines in Figure 1A mark the significance thresholds for a one-sample t-test at a = 0.001 for sample sizes *N* = 20 (gray lines) and *N* = 100 (black lines). Mirroring the positive whole brain means, more voxels with positive COPE means pass the significance threshold for all tasks. The gambling task COPE values are smaller than those observed in the other three tasks while their standard deviation is almost as large. This results in less significant voxels for the gambling task compared to the other tasks.

**Figure 1:**
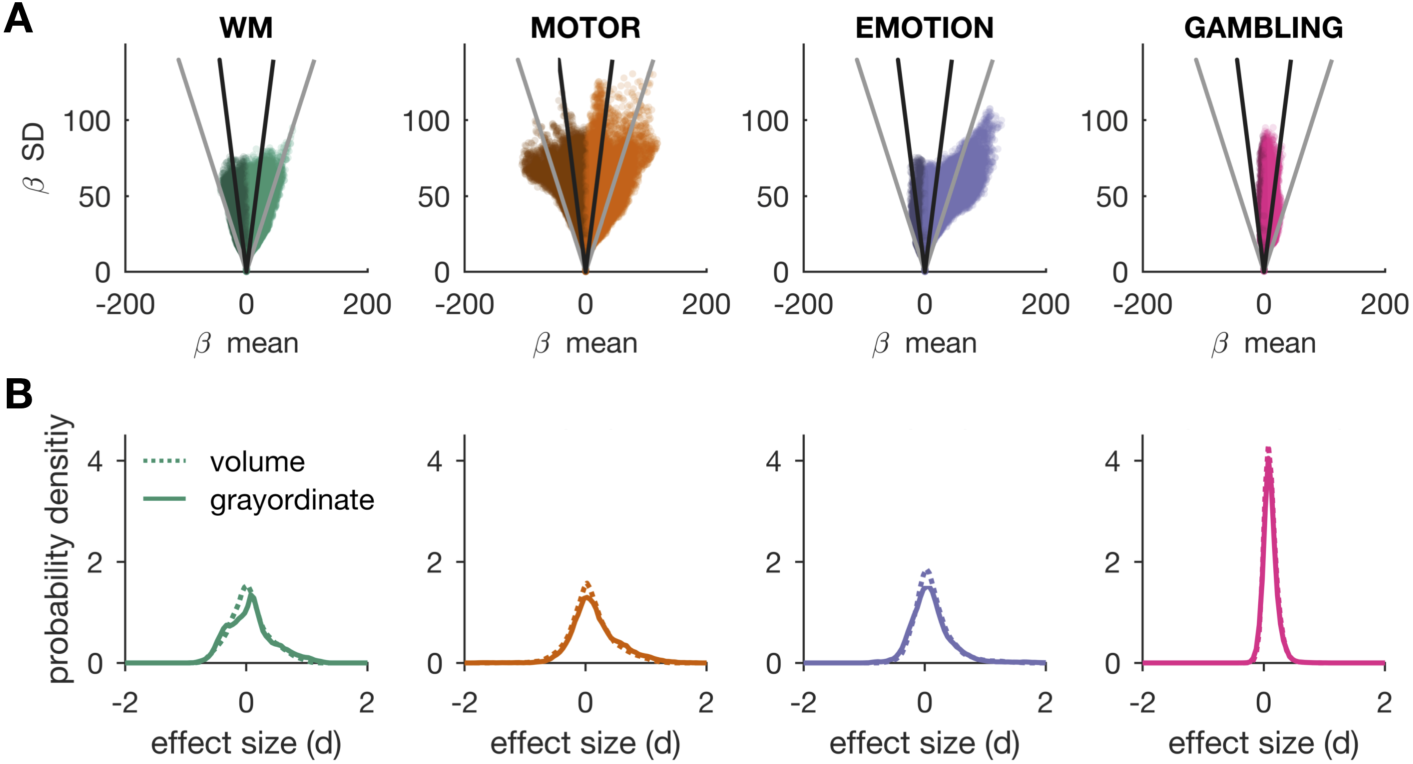
(A) The voxel-wise standard deviation vs. mean of the contrast estimates for working memory (WM), motor, emotion and gambling tasks. Voxels with positive and negative β means are depicted using different brightness. Voxels under “V”-shaped lines are significant for sample size *N* = 20 (gray) or *N* = 100 (black) and using a threshold of *p <* 0.001. (B) Probability density functions for effect sizes (Cohen’s d) for all tasks comparing volumetric (dotted lines) against grayordinate (solid lines) formats.

Figure 1B shows probability density distributions of the the population effect sizes for each of the four tasks for both volumetric (dotted lines) and grayordinate data (solid lines). Both data types are in good agreement. The grayordinate data are slightly shifted to the right for the motor task and slightly shifted to the left for the emotion task.

Table 1 shows the proportion of voxels (or vertices) that fall in each effect size category. Small effects are present in about 16% of the voxels for the gambling task, and about 30 ^_^ 38% for the other three tasks. Medium effects are present in about 6 - 13% of the voxels for WM, motor, and emotion tasks, while large effects are only present in about 4 - 8% of the voxels. These numbers are much lower for the gambling task (< 1% large effects). Interestingly, the grayordinate data have higher proportions of large effects for all tasks but the improvement is particularly large for the motor task. Differences between volumetric and grayordinate data are small for the medium and small effect size categories.

**Table 1:** Proportions of voxels (vertices) in percent for each effect size category organized by task and data format. Here ‘Vol’ represents volumetric and ‘Gray’ grayordinate.

|  | **WM** |  | **Motor** |  | **Emotion** |  | **Gambling** |  |
| --- | --- | --- | --- | --- | --- | --- | --- | --- |
|  | Vol | Gray | Vol | Gray | Vol | Gray | Vol | Gray |
| Small $d$ | 35.02 | 37.90 | 33.01 | 32.10 | 30.85 | 33.15 | 16.87 | 16.15 |
| Medium $d$ | 11.31 | 12.88 | 11.08 | 12.66 | 6.75 | 8.08 | 0.49 | 0.53 |
| Large $d$ | 2.06 | 4.78 | 4.70 | 7.61 | 3.26 | 3.54 | 0.02 | 0.07 |

Figure 2 maps effect size results onto the brain. Results for the cortical surfaces are shown in (A) and volumetric results are shown in (B). Please note that grayordinate (A) and volumetric (B) results may differ for the cortex but not for subcortical regions. Overall, WM and emotion are relatively even in positive and negative effects. Motor and gambling have predominantly positive regions and a few regions that show negative effects.

**Figure 2:**
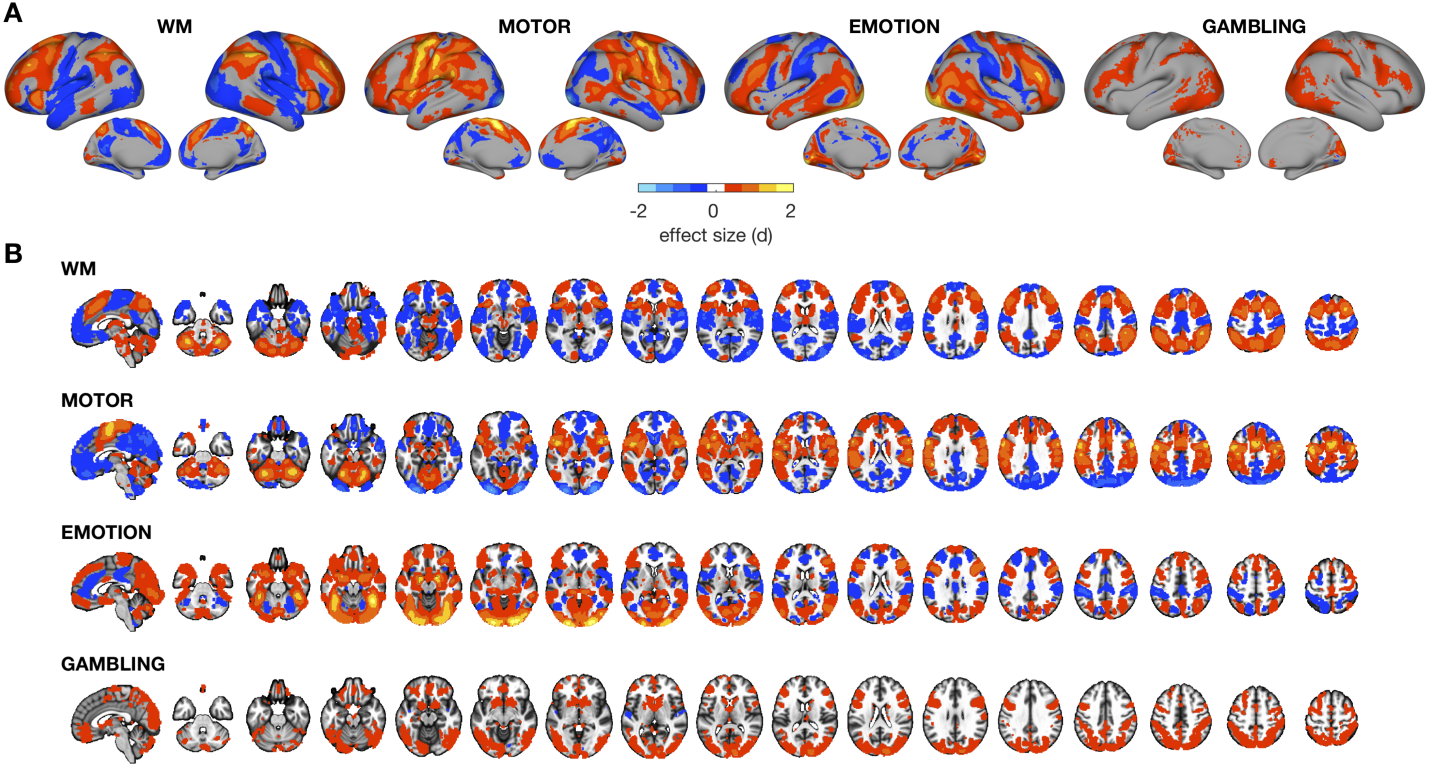
(A) Surface effect size maps for all tasks using the grayordinate data. (B) Effect size maps from the volumetric data. Please note that the data formats differ for the cortex but subcortical structures will be highly similar.

Large positive effects for the WM task are located in the dorsal lateral pre-frontal cortex (dlPFC), intraparietal sulcus (IPS), medial superior frontal gyrus (mSFG), and also in the cerebellum (Figure 2). Medium and small positive effects include most of the dorsal and lateral prefrontal cortex, parietal cortex, anterior insula, and anterior midcingulate cortex (aMCC). Negative effects are overall smaller and were observed in the regions often referred to as ‘task negative’. Negative effects are most pronounced in the retrosplenial cortex (RSC), ventrolateral prefrontal cortex (vlPFC), medial temporal lobe (MTL), and parietal operculum.

For the motor task, the largest effects are located in the motor cortex in the prefrontal sulcus (M1), supplementary motor area (SMA), and cerebellum. Other large positive effects were observed in sensory regions in the postcentral sulcus (S1), the parietal operculum, and the supramarginal gyrus. Negative effects are more sparse with the strongest negative effects located in the occipital cortex.

The strongest positive effects for the emotion task are located in the occipital cortex, the fusiform gyrus, amygdala, and the right lateral PFC. Negative effects are located in the dorsal parietal cortex and in multiple smaller regions on the medial walls.

Effects in the gambling task are smaller than in the other tasks. Positive effects are located in the lateral PFC, cerebellum, occipital and parietal cortices. Positive but small effects were also observed in the ventral striatum and basal ganglia.

Figure 3 shows the sample size required to achieve a certain level of power. As the sample size increases, the power of the t-test increases assuming a fixed effect size. This is visualized by the inverse relationship between effect size in Figure 2 and the sample size needed in Figure 3. In most cases, the biggest jump in power occurs between *N* = 20 and *N* = 40, with a steady increase when *N* > 40. This suggests that *N* = 40 may be a good sample size for individual fMRI studies when taking into consideration both reliability and cost. For the robust tasks analyzed here, even sample sizes of *N* < 10 theoretically offer 50% power for the WM, motor, and emotion tasks. For example, M1 and SMA in the motor task and IPS in the WM task achieve 50% power with such small samples, which can help explain the enormous success of small sample fMRI studies in the early 1990’s relying primarily on such robust tasks. Sample sizes of *N* = 20 to *N* = 40 achieve 80% power in primary sensory and motor areas for the main effects of motor activity or working memory load. Again, the gambling task stands out in that much larger sample sizes are needed to obtain reasonable statistical power. Some occipital regions achieve 80% power with samples of *N* > 20 while samples of *N* > 40 achieve 50% power in the ventral striatum. Since the ventral striatum is a key region for value learning that has been consistently associated with the processing of unexpected rewards (Knutson et al., 2001; Gläscher et al., 2010; Haber and Knutson, 2010), the low power observed here is likely related to choice of a block design for the subject-level models. Power maps for the volumetric data are shown in Supplementary Figures 8-11. Results for both data types are again very similar.

**Figure 3:**
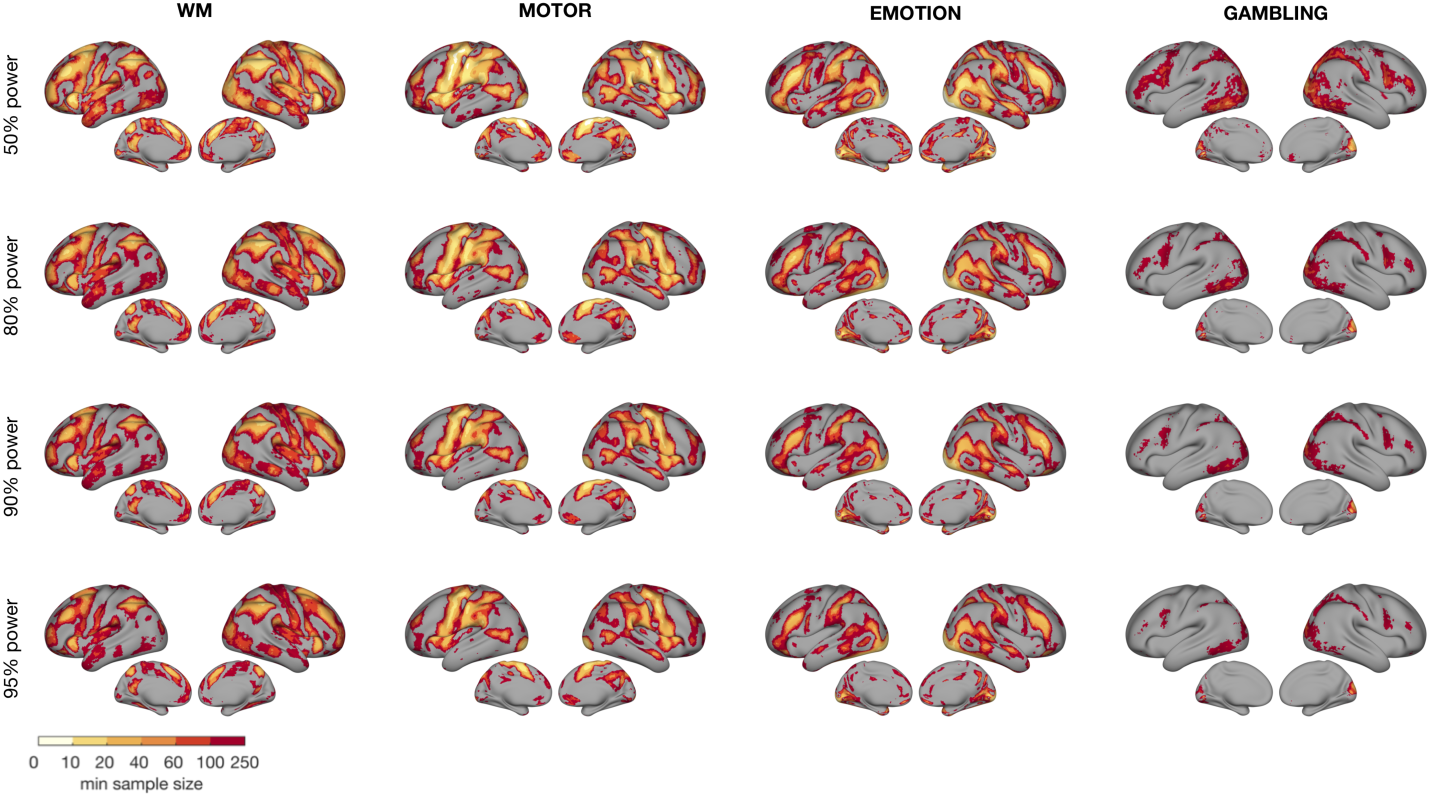
Power maps for surface data for all tasks. The minimum sample sizes required to achieve 50, 80, 90, and 95% power are indicated by different colors.

### 3.2 Effect of Sample Size on Group Analysis

Figure 4 shows the proportion of times each voxel is significant in the *K* repetitions for each sample size and task. The empirically determined power maps are in agreement with the theoretically derived minimum sample size maps shown in Figure 3 and the population effect size maps in Figure 2. Furthermore, volumetric (Supplementary Figures 12-15) and grayordinate results largely converged. For larger effect sizes in the WM, motor, and emotion tasks, empirical power quickly jumps from 50 to 95% and more. This sharp increase in empirical power is particularly prominent in samples of *N* = 60 and larger in the lower half of Figure 4. Even for extremely small samples of *N* = 10, a number of voxels in the motor and emotion tasks achieve 80% power while a few voxels in the WM task reach at least 50% power. The voxels with the largest effects in the WM, motor, and emotion tasks already achieve 95% power with sample sizes of *N* = 20-40. Even a few occipital voxels in the gambling task reach 95% power at *N* = 40. These empirical power estimates show that with very robust tasks at least some significant results are almost guaranteed with sample sizes of *N* = 20. However, the qualitative differences between in power maps for 20 and 80 subjects show that smaller studies may miss many of the wide-spread medium-sized effects, for example lateral prefrontal, temporal-parietal, and retro-splenial cortex in the motor task.

**Figure 4:**
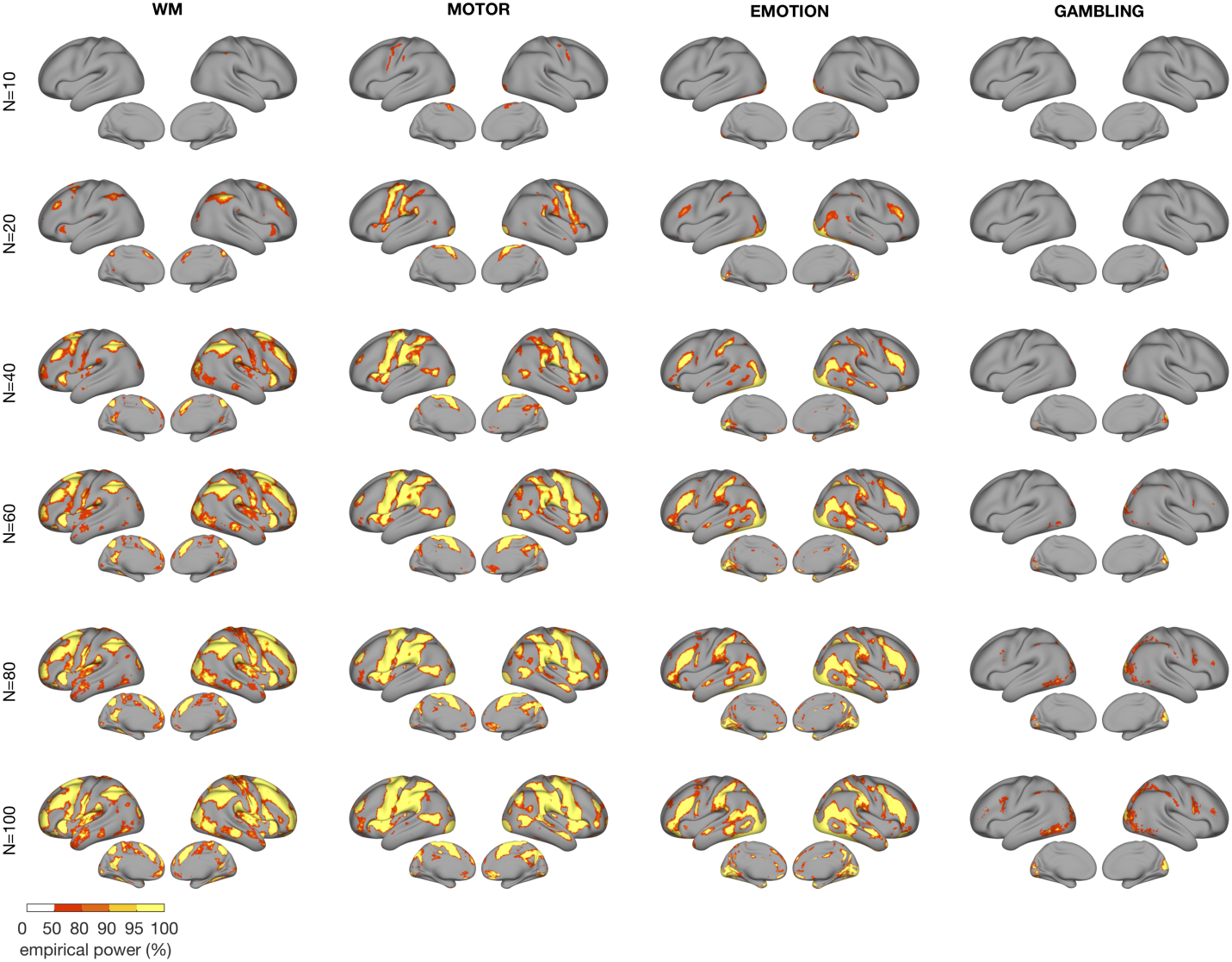
Empirical power maps for the surface data. Empirical power is estimated as the fraction of *K* = 100 repetitions of sample size *N* in which any given voxel was significant at *p <* 0.001.

Next, we computed summary statistics for the each effect size category (i.e., small, medium, and large effects). For each category, we computed the proportion of voxels that are deemed significant in at least 80% of samples when thresholding at *p <* 0.001 (Figure 5). The highest proportion of active voxels in at least 80% of samples was observed in the emotion task. Here, 6.8% of the large effect voxels were active in 80% of the samples with *N* = 10, or in other words, achieved 80% empirical power. This indicates that the vast majority of large effects will be missed with 10 subjects, although the empirical power maps in Figure 4 suggest that a few voxels will pass the significance threshold. The proportion of active voxels increases steeply for larger sample sizes in all tasks for the large effect sizes. For the medium sized effects, the curves are shifted to the right, indicating the necessity of larger samples, as expected. With 40-60 subjects per sample, the proportion of voxels with at least 80% empirical power exceeds 70% of the voxels in the large and medium categories (except for the grayordinate gambling task data). Samples of 80 subjects are sufficient to detect almost all large and medium effects. By contrast, even sample sizes of *N* = 100 are very unlikely to detect any small effects reliably as indicated by the low proportions active for all tasks. The volumetric and grayordinate data formats differ little in this analysis. The volumetric format fares slightly better at detecting medium effect sizes at small to medium sample sizes.

**Figure 5:**
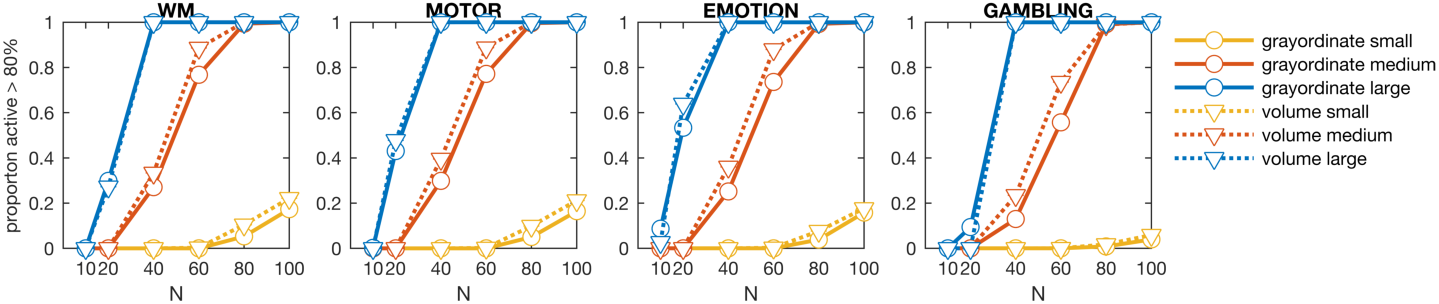
Proportion of active voxels. The proportion of voxels in each effect size category (indicated by colors and symbols; see legend) that are deemed significant in at least 80% of samples using a per-voxel threshold of p < 0.001 for each of the four tasks. Results are shown for grayordinate (solid lines) and volumetric (dotted lines) data.

The previous analyses asked which effects can be recovered at different sample sizes. The next analyses investigated which effects might be missed at different sample sizes. Figure 6 shows the average proportion of voxels in each effect size group that are not deemed active (i.e., are considered false negatives under the assumption that voxels in that category should be active) when thresholding at *p <* 0.001. For the high effect-size voxels, a sample consisting of 40 subjects is sufficient to reliably detect each voxel. For the medium effect-size voxels, a sample consisting of 80 subjects is sufficient to reliably detect each voxel. To reliably detect voxels in the low effect-sizes group, samples of well over 100 subjects are required. Interestingly, for *N* = 20 there are between 20 - 40% false negatives for the high effect size group depending on the task. For the medium effect sizes, samples of 40 subjects achieve less than 40% false negatives and samples of 80 - 100 subjects result in almost no false negatives. Again, differences between volumetric and surface base data formats are small, with the volumetric data offering slight improvements over the grayordinate data.

**Figure 6:**
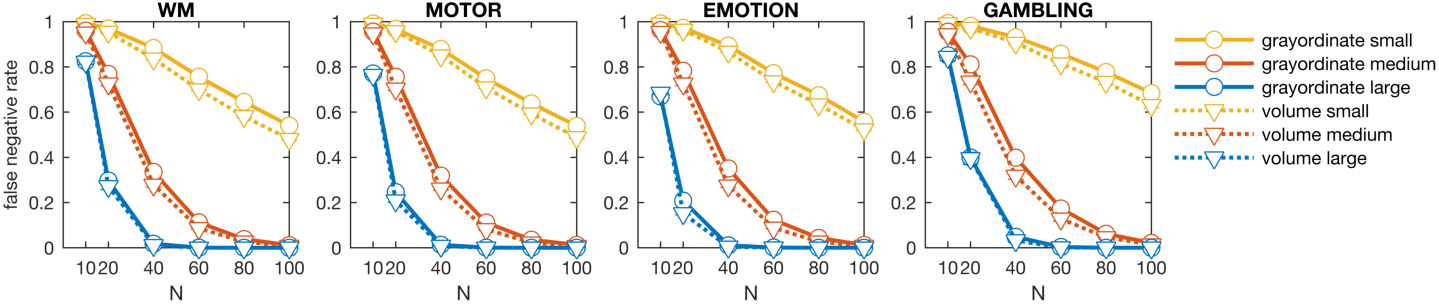
False negative rates. The average proportion of voxels in each effect size category that were not deemed active (i.e. are considered false negatives under the assumption that voxels in that category should be active) when thresholding at p < 0.001. Results are shown for grayordinate (solid lines) and volumetric (dotted lines) data.

### 3.3 Effects of Circular Analysis on Effect Size Estimation

To estimate how much effect size estimates from small samples overestimate the true effect size in non-independent analyses, we compute the absolute difference between sample size effect sizes and the population effect size for the same cluster. For each significant cluster at threshold p < 0.001, we compute the average effect size and plot the difference between this value and the population effect size; see Figure 7 for results for grayordinate data. The volumetric results are highly similar and shown in Supplementary Figure 16. For small samples of 10 subjects effect sizes are inflated by 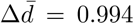 for the motor task to 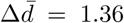 for the gambling task. The effect size bias decreases for larger samples but the mean bias still 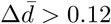 for samples with 100 subjects. Using an completely independent region-of-interest definition, for example based on an atlas (Glasser et al., 2016; Shen et al., 2013), mitigates this effect size inflation (maximum mean bias 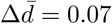 for *N* = 10 and less for larger samples).

**Figure 7:**
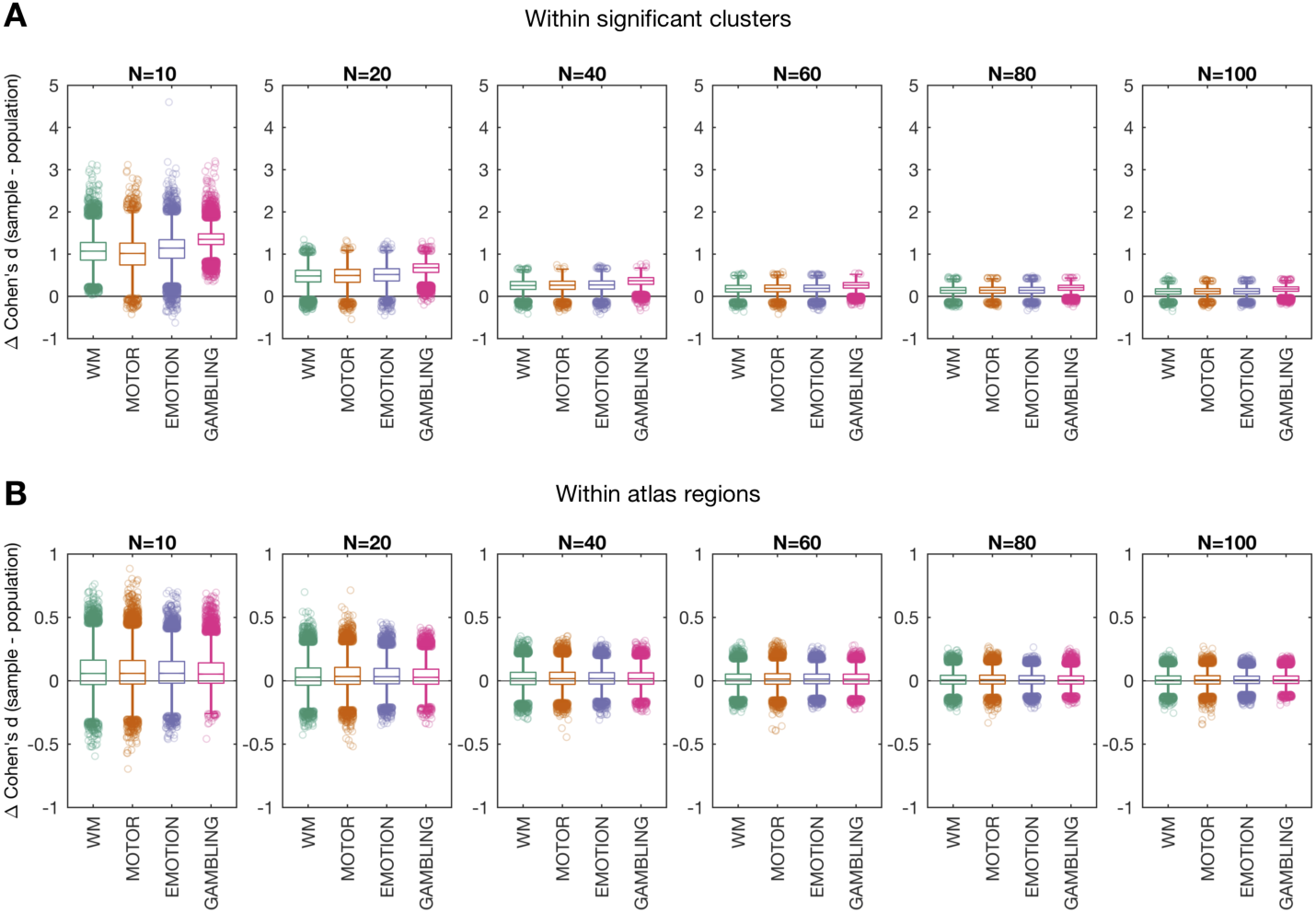
Effect size bias. (A) For each sample of size *N* the empirical effect size *d* was computed based on its average within each significant cluster. The population effect size for that cluster was then subtracted from this value. Boxplots show the distributions across all clusters from all 100 samples for each task and sample size. (B) same plot showing the differences between the average sample effect sizes and the population effect sizes, now for each parcel from the atlas of Glasser et al. (2016).

## 4 Discussion

In this work, we study the effects of sample size on the sensitivity and reliability of group-level OLS analysis. Treating the complete collection of subjects (*n* = 491) from the 500-subject release from the Human Connectome Project (HCP) as our population of interest, we evaluated the ability of small sampled studies to effectively capture population-level effects of interest. Our study shows that sample sizes of 40 are generally able to detect regions with high effect sizes (Cohen’s *d >* 0.8), while sample sizes closer to 80 are required to reliably recover regions with medium effect sizes (0.5 < d < 0.8). In general, we show there is low sensitivity to detect small effects (d < 0.5) for any reasonable fMRI sample size.

The analysis was repeated both for volumetric data and surface based data. The results were not noticeably different between the two data types, indicating that working on the surface does not provide improvements in terms of sensitivity when performing a massive univariate GLM analysis. The surface based data had a higher proportion of large effects but differed less with respect to medium effect sizes. This would suggest that surface based analyses might be more sensitive for smaller samples. However, this was not observed for the proportion of voxels deemed active or false negative rates. It remains possible that surface based analysis will be preferable if a spatial model is used instead; see, for example Mejia et al. (2017). However, this topic is outside the scope of the current manuscript.

This work can be seen as an expansion of the work by Thirion et al. (2007), who used a cohort of 80 subjects to investigate the role inter-subject variability plays in sensitivity and reliability of studies with a smaller number (e.g., 10 - 16) of subjects. They showed, among other things, that 20 subjects or more should be included in functional neuroimaging studies in order to have sufficient reliability. Since our analysis was based on a larger data set, this allowed us to investigate larger sample sizes, more variable tasks, and data sampled at a higher spatial and temporal resolution than possible in Thirion et al. (2007). Our findings suggest a recommend sample size of 40 or more subjects. A notable exception are areas such as M1 and SMA in the motor task and IPS in the WM task, which achieve 50% power with samples around 10. This no doubt explains the success of early small sample fMRI studies that relied primarily on such robust tasks.

The HCP task fMRI data set we used has appeared in several recent papers to investigate related topics. For example, in Poldrack et al. (2017) the data was used to obtain effect-size estimates for a number of common neuroimaging experimental paradigms in certain regions of interest. Using a smaller cohort of subjects (*N* = 186) they showed that effect sizes in fMRI are surprisingly small, even for powerful tasks such as the motor task. They conclude that the average fMRI study is poorly powered for capturing realistic effects. In Lohmann et al. (2017) the data was used to investigate the false negative rates in studies with small sample sizes. Similar to our approach, they used a larger cohort of subjects (*N* = 400) to approximate the true positive effects. Thereafter, they computed the fraction of those effects that was detected reliably using standard software packages at various smaller sample sizes. They found that for common sample sizes this value was generally below 25%. In this work, we validate and expand upon the results from these papers.

It is interesting to note that samples well exceeding 100 subjects are needed to reliably detect voxels with effect sizes ranging between 0.2 and 0.5. Thus, using sample sizes available to most individual labs we appear far from the situation warned about in Friston (2012) that large sample sizes produce statistically significant effects of no practical importance (i.e. ‘trivial effects’); see Lindquist et al. (2013) for a rebuttal of this point. Friston (2012) presented an analysis of effect size in classical inference that suggested the optimal sample size for a study is between 16 and 32 subjects. For larger studies even the smallest treatment effect will appear significant. In this study we show that this does not appear to be a realistic concern, particularly given that the HCP data are of unusually high quality and effect size estimates are likely higher than those that would be found in most standard fMRI studies.

In order to estimate effect sizes for a planned study or grant, researchers often conduct a small pilot study. Even though the inflation of effect size estimates in circular analyses, i.e., computing effect sizes in significant clusters of the same or other non-independent COPEs (Kriegeskorte et al., 2010; Vul et al., 2009), is widely acknowledged, it is unclear how large this bias is in real data. Here, we show that non-independent analyses drastically overestimated effect sizes on average by 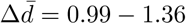, depending on the task when *N* = 10. Thus there is a risk that in computing effect sizes from pilot studies, researchers may be vastly overestimating their values and thus vastly underestimating the number of subjects required in the study. In general, we recommend that researchers use masks defined by the intersection between functional activation and anatomical masks, as discussed in Poldrack et al. (2017), to perform effect size estimation.

Results for the gambling task stood out from the other tasks. Overall, effect sizes in the gambling task COPE were much lower than for the other tasks, leading to less statistical power and requiring larger samples to reliably detect brain activation, even in the ventral striatum, a key brain region for reward learning (Knutson et al., 2001; Gläscher et al., 2010; Haber and Knutson, 2010). The relatively small effect sizes for the gambling COPE may be caused by modeling multiple reward and loss events as blocks at the subject level instead of using event-related regressors (Barch et al., 2013). While a more detailed evaluation of this particular COPE is beyond the scope of this study, future studies may consider investigating the effects of the short task duration, or the low regional SNR (Barch et al., 2013) for this task in more detail.

Overall, our results support the current movement towards conducting larger fMRI studies. Acquiring at least 40 subjects for within-group hypothesis about regional activations or deactivations allows for reasonable power to detect large effects. Acquiring even larger and more costly samples, will enable researchers to investigate the functions of brain regions with medium sized effects, which may cover about 30% of the graymatter voxels.

## Acknowledgements

Data were provided by the Human Connectome Project, WU-Minn Consortium (Principal Investigators: David Van Essen and Kamil Ugurbil; 1U54MH091657), which was funded by the McDonnell Center for Systems Neuroscience at Washington University and the 16 NIH Institutes and Centers that support the NIH Blueprint for Neuroscience Research. The work presented in this paper was supported in part by NIH grants R01 EB016061 and P41 EB015909 from the National Institute of Biomedical Imaging and Bioengineering and by the DFG (GE 2774/1-1).

## Footnotes

1 https://wiki.humanconnectome.org/pages/viewpage.action?pageId=88901591

